# Reduced Survival Expectancy Weakens Reproductive Isolation Between Annual Fishes *Garcialebias reicherti* and *G. charrua*

**DOI:** 10.1101/2023.12.06.570411

**Authors:** Noelle Rivas-Ortiz, Carlos Passos

**Affiliations:** Sección Etología, Facultad de Ciencias, Universidad de la República, Montevideo, Uruguay

**Keywords:** Aggression, annual fishes, hybridization, mate choice, reproductive effort, sexual selection, survival expectancy

## Abstract

Hybridization depends on reproductive isolation, which can be impacted by mate choice. Mate choice may vary temporally, as it is modulated by several factors, including survival expectancy and future opportunities for reproduction. *Garcialebias reicherti* and *G. charrua* are annual fishes with parapatric distributions that hybridize in the overlapping area of their distributions. They inhabit temporary ponds that flood during the autumn and dry out during the spring, resulting in decreased survival expectancy and future opportunities for reproduction during the breeding season. We predicted that a decrease in survival expectancy would promote reproduction and reduce reproductive isolation between *G. reicherti* and *G charrua*. By simulating desiccation in the early and late breeding season, we investigated the effects of the desiccation risk and the phase of the breeding season on reproductive isolation and reproductive effort of these species. As expected, our findings reveal that decreased survival expectancy influences both reproductive isolation between *G. reicherti* and G*. charrua*, and their reproductive effort. Notably, reproductive isolation between these species decreased under a high desiccation risk and in the late breeding season. Additionally, we observed an increase in the frequency of mating and courtship events and aggressive behaviours in the late breeding season. Our study suggests that reproductive isolation between *G. reicherti* and *G. charrua* and their reproductive effort can change rapidly within a short period of time, emphasizing the influence of survival expectancy on the temporal dynamics of reproductive isolation and hybridization.

## INTRODUCTION

Hybridization in animals was traditionally considered an unusual event. However, it is currently known that individuals of different species can reproduce with each other and that it is prevalent in some natural populations (Abbott et al., 2013; Taylor & Larson, 2019). Furthermore, hybridization can occur over long periods of time (Mallet, 2005), and it can even reach stable equilibria in which species maintain their identities despite gene flow (Irwin & Schluter, 2022; Servedio & Hermisson, 2020). Most research on hybridization has examined its geographic variation (Sato et al., 2016; Yang et al., 2016; Yukilevich & Peterson, 2019) or its long-term changes (e.g. Kulmuni et al., 2020; Seehausen et al., 2014). Here, we examine how changes in sexual behaviour can drive changes in hybridization in shorter time-scales.

Hybridization occurs in several contexts and depends on the evolutionary history and ecology of the species as well as behavioural decisions of individuals (Rosenthal, 2013, 2017). Hybridizing propensity depends on the degree of reproductive isolation between species, which can be strongly affected by sexual selection. To maximize fitness, selection is expected to favour mechanisms that reduce the probability of hybridizing early in the mating process (Rosenthal, 2017). Therefore, mate choice plays a major role in hybridization, as it can maintain reproductive isolation by reducing heterospecific mating (e.g. Atsumi et al., 2019; Pauers & Grudnowski, 2022; Shahandeh et al., 2018). Environmental and ecological factors often impact reproductive preferences (see Ah-King, 2019; Ah-King & Gowaty, 2016; Jennions & Petrie, 1997; Miller & Svensson, 2014) and thereby can also impact reproductive isolation (Rosenthal, 2013, 2017). In fact, it has been proposed that when choosiness is reduced in response to some environmental factor without a decrease in motivation to mate, hybridization could increase (Rosenthal, 2017). Survival expectancy, which also indicates future opportunities for reproduction, can affect both choosiness and reproductive effort. When survival expectancy is reduced, individuals are predicted to decrease their choosiness (Gowaty & Hubbell, 2005, 2009; Henshaw, 2018; Watts et al., 2022), a prediction supported by several experimental studies (Edomwande & Barbosa, 2020; Pilakouta et al., 2017; Willis et al., 2012). Similarly, individuals of some species have no preference for mates of their same species, or they even prefer heterospecific partners (Pilakouta et al., 2017; Rosenfield & Kodric-Brown, 2003; Sato et al., 2016; Wyman et al., 2014), when time constraints for breeding (Lynch et al., 2005; Nuechterlein & Buitron, 1998; Pfennig, 2007), or scarcity of mates (Izzo & Gray, 2011; Nuechterlein & Buitron, 1998; Willis et al., 2011) threaten their future reproductive opportunities. Hybridization in these situations could be adaptive, either because it is better to mate with a heterospecific than a conspecific or because it is better to mate with a heterospecific than not at all (Pfennig, 2021; Willis, 2013). Additionally, survival expectancy can also impact reproductive effort. Because resources are limited, individuals tend to face a trade-off between reproduction and survival (Edward & Chapman, 2011; Stearns, 1989). Therefore, when survival expectancy is threatened, individuals could reduce their current reproduction in order to enhance their likelihood of survival and future reproduction (Duffield et al., 2017). However, when future reproductive potential is too low, it would be advantageous to promote reproductive investment rather than suppress it (Clutton-Brock, 1984; Duffield et al., 2017; Williams, 1966).

Here we examine reproductive isolation and reproductive effort of two species of fishes that annually face the threat of desiccation. Annual fishes inhabit temporary ponds that flood during autumn and completely dry out during spring, causing adults to die in the late breeding season. Females lay desiccation-resistant eggs that hatch when the ponds flood again the next year (Berois et al., 2016; Wourms, 1972). This transient environment has selected for an accelerated life history. Fish reach adulthood rapidly (Blažek et al., 2013) and then reproduce continuously until the inevitable desiccation of the pond (Wourms, 1972). *Garcialebias*, previously called *Austrolebias* and recently renamed (Alonso et al., 2023), is a genus of South American annual fishes. We studied sister species *Garcialebias reicherti* and G*. charrua,* that inhabit ponds in eastern Uruguay (Garcia et al., 2009) and are parapatrically distributed (Figure 1). These species hybridize in nature, where their distributions overlap (Garcia et al., 2020), and in the laboratory, where they can reproduce in every possible combination and produce viable hybrids (Passos et al unpublished data). They are highly sexually dimorphic. Females have brown bodies and transparent fins and are almost indistinguishable between species (Garcia et al., 2009), while males have a clear background with dark vertical bands that differ between species (Loureiro & Garcia, 2008). Sexual selection drives their sexual behaviour through mate choice and male competition. They engage in elaborate courtship in which the males perform complex displays (García et al., 2008; Passos et al., 2016). Females show preferences for larger mates and conspecific males (Blengini et al., 2018; Passos et al., 2013), and males display intense male-male competition, rapidly engaging in contests when they meet (Passos et al., 2013; Passos et al., 2016).

**Figure 1.**
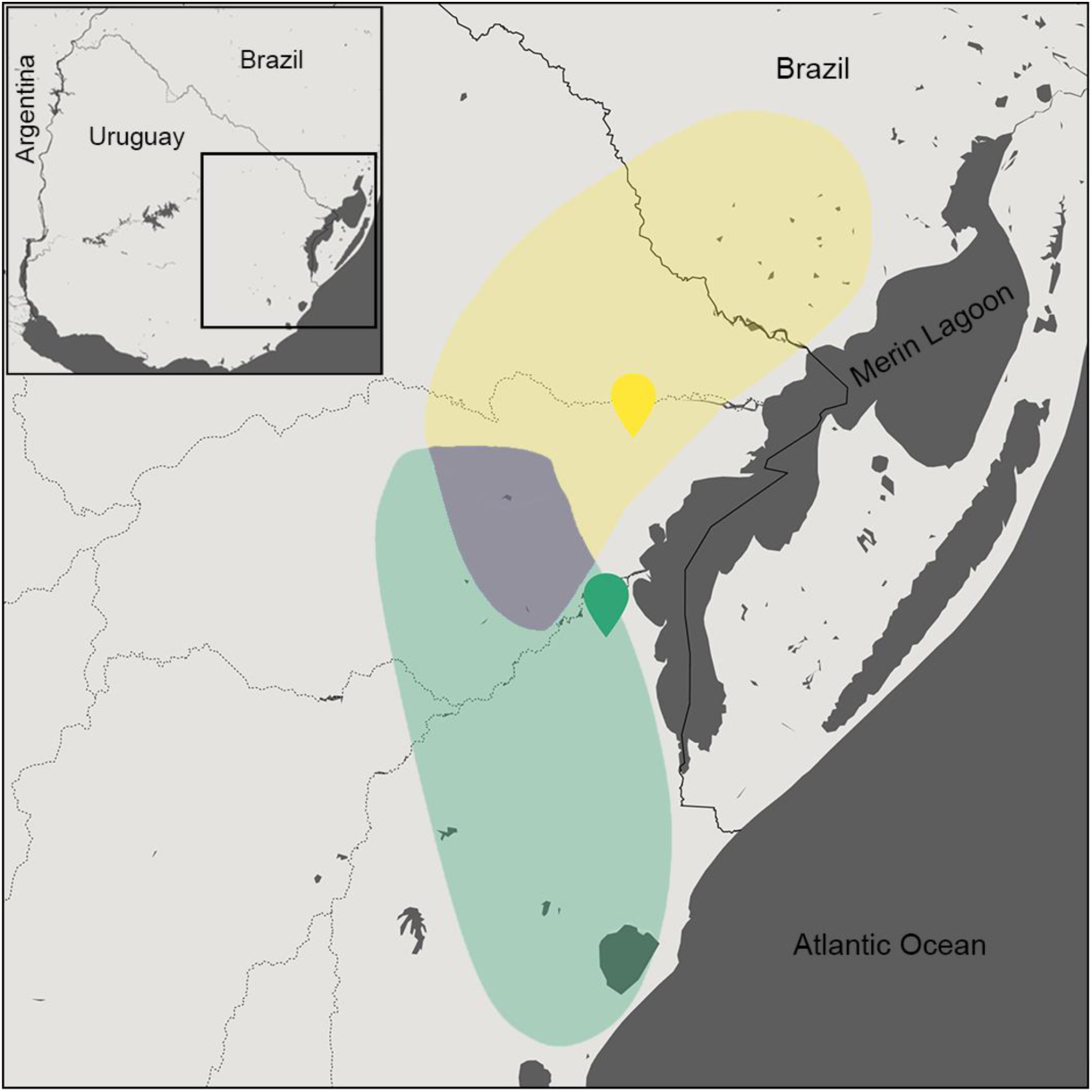
Distribution and collection sites of *Garcialebias reicherti* (yellow) and G*. charrua* (green). The violet zone is where their distributions overlap.

The duration of the reproductive period is quite restricted in *Garcialebias*. As the breeding season progresses and the temporary ponds dry out, survival expectancy and future reproduction opportunities decrease. *Garcialebias* have provided evidence of changes in their choosiness and reproductive investment while the breeding season progresses. Females, which prefer to mate with larger males in the early season, become less choosy in the late season (Passos et al., 2014). In addition, their reproductive investment also changes seasonally. Changes in gonadosomatic and hepatosomatic indexes of both females and males show that they allocate more energy to reproduction in the late breeding season, when both survival expectancy and future opportunities for reproduction decline (Passos et al., 2021).

In our study, we assessed whether the reduction in survival expectancy affected reproductive isolation between *Garcialebias reicherti* and *G. charrua*, and their reproductive effort. We evaluated the impact of both desiccation risk and the phase of the breeding season by experimentally simulating different desiccation risk scenarios in both early and late season. Then, we observed mating, courtship and aggression in behavioural assays. We predicted that under high risk of desiccation and in the late breeding season (1) reproductive isolation between *G. reicherti* y *G. charrua* would decrease, reducing assortative mating, and (2) their reproductive effort would increase, incrementing in the frequency of mating, courtship and male aggressive behaviours.

## METHODS

### Collection and Maintenance

Adult *Garcialebias reicherti* and *G. charrua* were collected from temporary ponds in the early and late breeding season (beginning of July and October 2021). G. *reicherti* was collected in Treinta y Tres Department (32°46’35.0”S, 53°39’38.0”W), and *G. charrua* was collected in Rocha Department (33°14’00.0”S, 53°44’03.5”W), Uruguay (Figure 1). Fish were kept in the laboratory under constant temperature (19° C) and natural photoperiod and fed daily with live *Tubifex sp*. They were housed in individual aquaria (20 x 8 x 16 cm, length × width × height), visually isolated from each other, for 36 h before experimental treatment.

### Desiccation Risk Experimental Treatment

We simulated different desiccation risk scenarios by housing groups of 12 individuals, three of each sex and species, in aquaria (57 x 37 x 20 cm) with different water depth for two weeks. Low desiccation risk housing aquaria (N = 4 for both early and late season) had a water depth of 15 cm, and high desiccation risk housing aquaria (N = 4 for both early and late season) had a water depth of 5 cm. All housing aquaria had two trays with peat moss as a spawning substrate and equal amounts of *Vesicularia dubyana* and stones, which served as refuges. Once a week, half of the water was replaced with clean filtered tap water.

### Behavioural Assays

Behavioural assays consisted of males and females interacting freely during three consecutive days. Each trial used one male and one female of each species (i.e., four individuals), held together in a test aquarium (40 x 13 x 16 cm) containing peat moss as spawning substrate. The test aquaria had the same water depth than the housing aquaria where the individuals lived (i.e., 15 cm in low desiccation risk housing aquaria, 5 cm in high desiccation risk housing aquaria). The left flank of all fish was photographed, and digital images were used to measure their standard length (from anterior of head to distal margin of caudal peduncle) using ImageJ version 1.53 (Schneider et al., 2012). Individuals of similar body size were chosen for each test aquarium, as body size is an important factor in both intrasexual and intersexual selection in *Garcialebias* (Passos et al., 2013, 2019).

We carried out seven trials for both high and low desiccation risk treatments in early breeding season, and six trials for both high and low desiccation risk treatments in late breeding season. We performed four daily observations, 10 min each, separated by 1 h intervals, recording conspecific and heterospecific mating and courtship events, and aggressive behaviours. Courtship is similar in both species, consisting of lateral displays in which the male extends its fins and quivers his body, and sigmoid displays, in which he performs body undulations (García et al., 2008; Passos et al., 2013). Mating events are unambiguous; the pair digs into the substrate and the male presses the female against it until she releases the egg (Passos et al., 2016). Male-male aggression is also similar in both species and consists of attacks, lateral displays, sigmoid displays and chases (Passos et al., 2016). Females of both species are visually indistinguishable (Garcia et al., 2009); therefore, we used individuals’ natural marks to identify them.

### Data Analysis

To assess the relationship between the recorded behaviours, the desiccation risk, and the phase of the breeding season, we used generalized linear mixed models (for mating and courtship) and generalized linear models (for aggression). Negative binomial distributions and log link functions were used in every model. For all models, we included the desiccation risk (high vs. low desiccation risk), the phase of the breeding season (early vs. late season) and the interaction between them as predictors (fixed effects). For mating and courtship models, we also included as predictors the mating or courtship type (conspecific vs. heterospecific) and its interaction with the other predictors. It was done to examine the differences between conspecific and heterospecific interactions, and their relationship with the desiccation risk and the phase of the breeding season. Courtship and mating models had trial identity as random effect, included as intercept, to account for repeated measures of each type of mating or courtship event. For all models, we conducted stepwise model simplifications and determined the predictors’ significance using LRT tests between nested models (Inchausti, 2023). Significance values were obtained by running ANOVAs using the car package, version 3.1.2 (Fox & Weisberg, 2018). When it was needed, post-hoc comparisons were performed by Tukey-adjusted test, using the emmeans package, version 1.8.5 (Lenth, 2023). All the considered models’ residuals were visually analysed using the DHARMa package, version 0.4.6 (Hartig, 2022), and they showed a satisfactory fit (results not shown). To be certain that there were no differences in the body size of individuals used in behavioural assays, we tested for differences in their body length. For data that were distributed normally, we performed t-tests for paired data, and for data that violated the assumption of normality, we applied Wilcoxon test. Data were checked for normality using Shapiro-Wilk test. Tests showed no significant differences in fish body size (p>0.05 for every test). Statistical analyses were performed using R version 4.3.0 (R Core Team, 2023) and the library glmmTMB version 1.1.7(Brooks et al., 2017). Differences were considered significant when P ≤ 0.05.

### Ethical Note

All experimental procedures complied with ASAP/ABS Guidelines for the Use of Animals in Research and were approved by our institutional ethical committee (approval number #240011-501204-21).

## RESULTS

### Reproductive Isolation

Even though males performed more conspecific than heterospecific courtship events, and it was not affected by the phase of the breeding season nor the desiccation risk (Figure 2, Table 1), assortative mating was impacted by both desiccation risk and the phase of the breeding season (Table 1). In the early breeding season, individuals from low desiccation risk trials performed significantly more conspecific mating events than heterospecific mating events (Tukey pairwise comparison, p < 0.001). By contrast, there was no difference between the number of conspecific and heterospecific mating events performed by individuals from high desiccation risk trials (Tukey pairwise comparison, p = 0.18) (Figure 3). In addition, in the late breeding season, there were no differences between the number of conspecific and heterospecific mating events in either high (Tukey pairwise comparison, p = 0.17) nor low (Tukey pairwise comparison, p = 0.27) desiccation risk trials (Figure 3).

**Figure 2.**
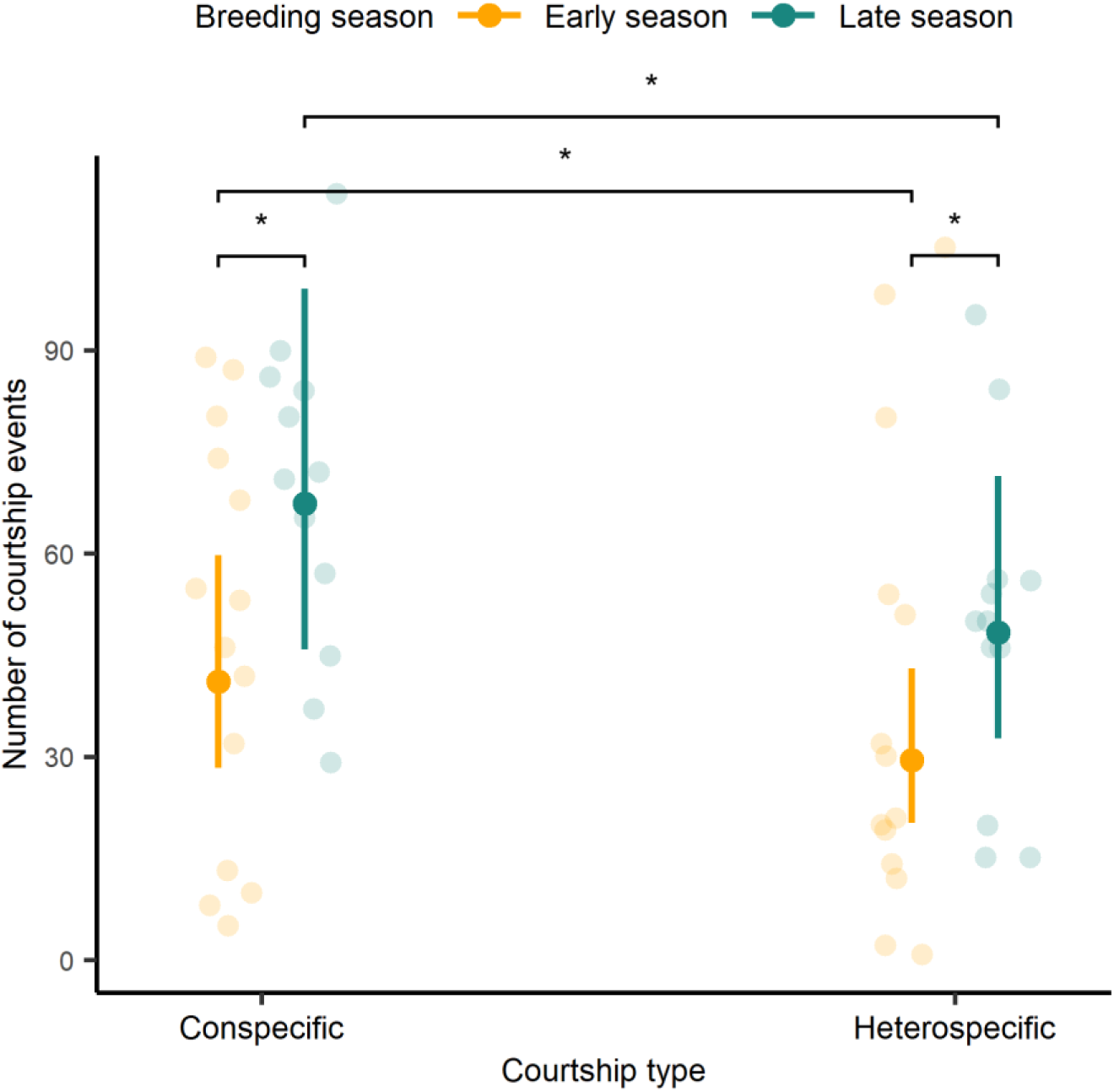
Effect of season on courtship events. Results are combined for both high and low desiccation risk trials, as the desiccation risk had no significant effect and was removed from the model. Orange points show courtship events performed in the early season, and green points show courtship events performed in the late season. The estimate means and their 95% confidence intervals are shown in solid-coloured points, and the raw data are show in more transparent points. (*) indicates p-value ≤ 0.05.

**Figure 3.**
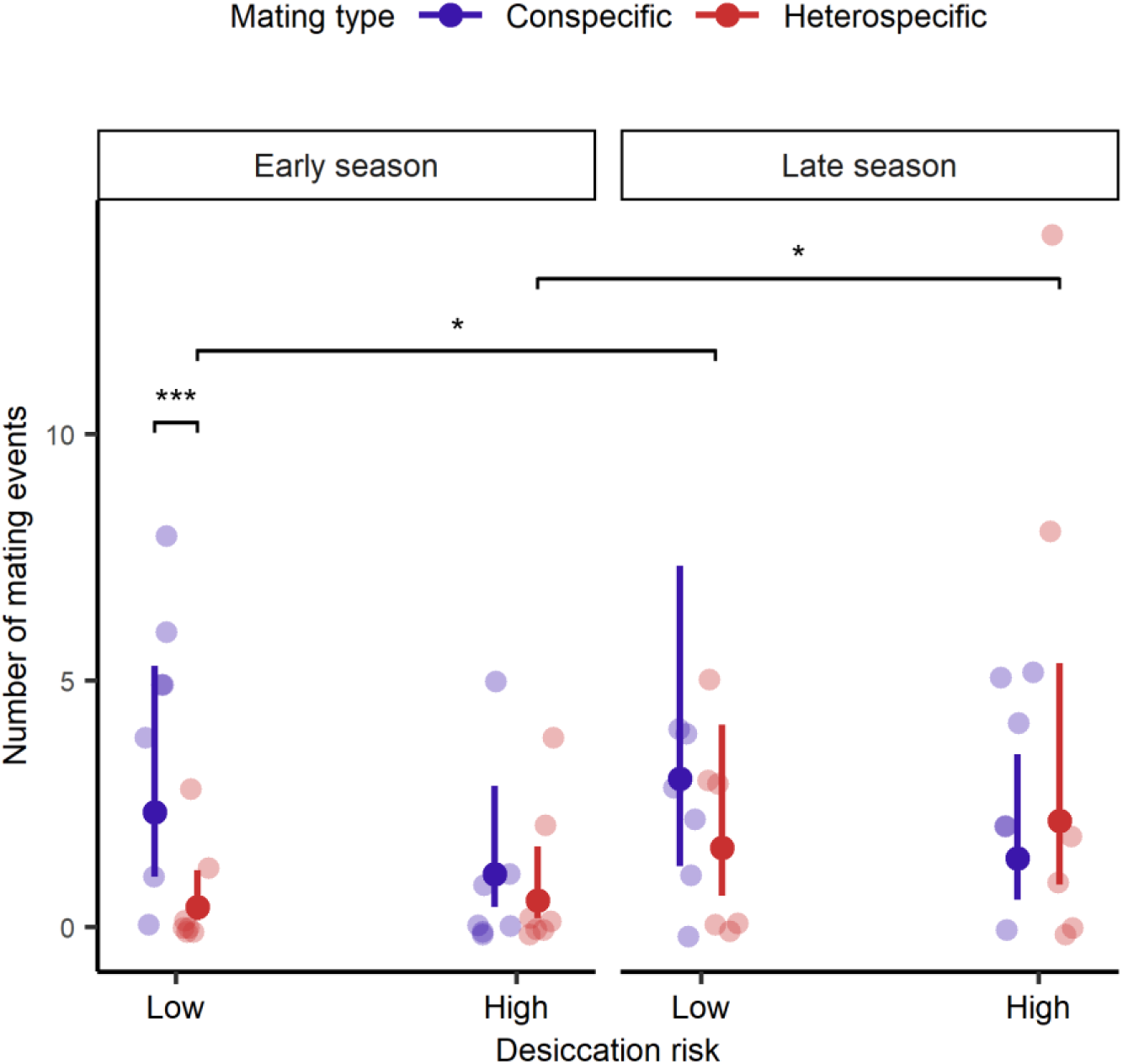
Effect of desiccation risk and breeding season on mating events. Blue points show conspecific mating events and red points show heterospecific mating events. The estimated means and their 95% confidence intervals are shown in solid-coloured points, while more transparent points show the raw data. (***) indicates p-value < 0.001, (*) indicates p-value ≤ 0.05.

**Table 1.** Impact of desiccation risk (high or low desiccation risk) and season (early or late season) on mating and courtship number and type (conspecific or heterospecific). Results of ANOVAs for the Generalized Linear Mixed Models. Both fixed and random effects are shown. For fixed effects, we show tests statistics (χ^2^), its degrees or freedom (between brackets) and p-values. For random effects, the standard deviation is shown. Significant p-values are highlighted in bold.

|  |  |  |  |
| --- | --- | --- | --- |
| Mating | Fixed effects | $\chi^2$(df) | p-value |
|  | Mating type | 3.08 (1) | 0.08 |
|  | Season | 2.04 (1) | 0.15 |
|  | Desiccation risk | 0.63 (1) | 0.43 |
|  | Mating type : Season | 4.43 (1) | **0.04** |
|  | Mating type : Desiccation risk | 4.23 (1) | **0.04** |
|  | Random effects | SD |  |
|  | Group identity | 0.93 |  |
| Courtship | Fixed effects | $\chi^2$(df) | p-value |
|  | Courtship type | 5.20 (1) | **0.02** |
|  | Season | 3.80 (1) | **0.05** |
|  | Random effects | SD |  |
|  | Group identity | 0.52 |  |

### Reproductive Effort

The frequency of mating events, courtship events and aggressive behaviours increased as the breeding season progressed. The number of heterospecific mating events was significantly higher in the late season than in the early season (Tukey pairwise comparison, p = 0.02). Conspecific mating events also increased in the late season, however, the difference between early and late season was not significant (Tukey pairwise comparison, p = 0.61) (Figure 3). In addition, males performed more courtship events in the late season than in the early season (Figure 2, Table 1), and the number of all aggressive behaviours (except attacks) significantly increased in the late breeding season (Table 2).

**Table 2.** Effect of season in aggressive behaviours. Results of the Generalized Linear Models and their ANOVAs. We show estimates, representing the predicted mean values of each behaviour in data scale, and their 95% confident intervals. We also show ANOVAs tests statistics (χ^2^), its degrees or freedom (between brackets) and p-values. Results are for both low and high desiccation risk, since desiccation risk had a non-significant effect and was removed from the models. Significant p-values are highlighted in bold.

| | | Estimate | 95% CI | $\chi^2$(df) | p-value |
| --- | --- | --- | --- | --- | --- |
| Attacks | Early season | 39.86 | 24.35, 65.23 | 0.04 (1) | 0.85 |
|  | Late season | 42.83 | 25.17, 72.88 |  |  |
| Lateral displays | Early season | 3.86 | 2.35, 6.32 | 8.09 (1) | **0.005** |
|  | Late season | 10.5 | 6.48, 17.00 |  |  |
| Sigmoid displays | Early season | 8.93 | 5.20, 15.32 | 9.43 (1) | **0.002** |
|  | Late season | 30.25 | 17.26, 53.02 |  |  |
| Chases | Early season | 23.64 | 15.46, 36.15 | 6.31 (1) | **0.01** |
|  | Late season | 52.25 | 33.30, 81.97 |  |  |

Desiccation risk did not affect the frequency of sexual behaviours. For both courtship and aggressive behaviours, desiccation risk was removed from the models (Tables 1, 2). For the mating events, it was retained due to its interaction with the mating type (conspecific vs. heterospecific), although it did not affect the number of mating events. There was no difference in the number of mating events performed by low and high desiccation risk fish, either for conspecific (Tukey pairwise comparison, p = 0.13) or heterospecific (Tukey pairwise comparison, p = 0.61) mating events, in any phase of the breeding season.

## DISCUSSION

In this study, we investigated whether reproductive isolation between *Garcialebias reicherti* and *G. charrua,* and their reproductive effort, varied according to survival expectancy. By simulating different desiccation risk scenarios in the early and late breeding season, we evaluated the effect of survival expectancy on assortative mating and on the frequency of mating, courtship and male aggressive behaviours. As expected, we observed that reproductive isolation decreased under a high risk of desiccation and in the late breeding season, and that reproductive effort increased in the late breeding season. Surprisingly, desiccation risk did not impact reproductive effort.

When survival is threatened, individuals tend to reduce their choosiness (Dougherty, 2021; Edomwande & Barbosa, 2020; Pilakouta et al., 2017), sometimes impacting assortative mating and reproductive isolation between species (Willis et al., 2012). Our results support our prediction that reproductive isolation between G*. reicherti* and G*. charrua* decreases under reduced survival expectancy. Even though there were more conspecific than heterospecific courtship events in both early and late season, reproductive isolation was weaker in the late season. While in the early season fish engaged in more conspecific than heterospecific mating events, in the late season there was no difference between them. Our results are consistent with previous evidence showing that *Garcialebias* females decrease their choosiness late in the breeding season (Passos et al., 2014), something that has also been found in other species (Borg et al., 2006; Nuechterlein & Buitron, 1998). While *Garcialebias* females prefer larger males early in the breeding season, they lose their preference in the late breeding season, when survival expectancy decreases (Passos et al., 2014). In addition to the breeding season, desiccation risk also affected the degree of reproductive isolation. In low desiccation risk trials, fish performed more conspecific than heterospecific mating events, however, in high desiccation risk trials there was no difference between the number of conspecific and heterospecific mating events performed. Thus, reproductive isolation was weaker under a high desiccation risk. These results are consistent with evidence found in *Spea bombifrons.* This amphibian also relies on temporary ponds for reproduction and its females increase hybridization when water depth is reduced (Pfennig, 2007). The impact of desiccation risk was evident only in the early breeding season, which could be because, in the late season, fish choosiness was already low due to the influence of the breeding season. Because the breeding season will inevitably come to an end, the progress of the breeding season might be a cue powerful enough to decrease choosiness independently of water depth.

When individuals have high chances of future reproduction, they are expected to reproduce at sub-optimal rates, in order to optimize the trade-off between current reproduction, somatic maintenance and future reproduction. However, when they have low chances of future reproduction, the opposite should happen. Individuals should increase their current reproductive investment in an effort to increase their offspring before death, even at a cost to their survival (Clutton-Brock, 1984; Duffield et al., 2017; Williams, 1966). This strategy, called terminal investment, has been documented in a great number of cases (e.g. Corbel & Carazo, 2022; Foo et al., 2023; Rutkowski et al., 2023). Our results parallel this evidence and follow our prediction; *G. reicherti* and *G. charrua* increase their sexual behaviour when survival expectancy is reduced over the breeding season. Fish performed more mating and courtship events in the late season than in the early season. In addition, male aggressive behaviours also increased in the late season, with more lateral displays, sigmoid displays and chases occurring the late breeding season than in the early breeding season. This behavioural change is in accordance with previous physiological evidence suggesting that *Garcialebias* increase their reproductive investment at the end of the breeding season (Passos et al., 2021). The seasonal changes in reproductive investment may be regulated by glucocorticoid hormones, such as cortisol (Passos et al., 2021). In *G. reicherti* natural populations, cortisol levels rise in the late season, and experimental cortisol treatment that simulates this natural cortisol rise, increases the number of courtship behaviours performed by males and intensifies their body coloration (Passos et al., 2021). In addition, the experimental cortisol treatment also changes fish body condition and gonadosomatic index, suggesting an increase in reproductive investment (Passos et al., 2021).

In contrast to the phase of the breeding season, and contrary to our prediction, desiccation risk did not impact the frequency of mating, courtship or aggressive behaviours. It could be because desiccation risk may have not impacted cortisol levels. Our experimental treatment might not have been appropriate to increase the level of cortisol or the expression of its receptors. The form and intensity of environmental cues that inform the survival expectancy are expected to indicate when to increase reproductive investment (Duffield et al., 2017). In that case, cortisol rise may require a combination of environmental cues, as in the ponds that *Garcialebias* inhabit, changes in water depth are usually accompanied by other environmental changes (Podrabsky et al 2016).

*Garcialebias* responded to a reduced survival expectancy over the breeding season by reducing their choosiness and by increasing their sexual behaviour. However, their response to an increased risk of desiccation was solely the reduction in reproductive isolation, as they did not increase their sexual behaviour. Different reasons could explain it. First, changes in choosiness and reproductive investment may have different costs and requirements. While the reduction in choosiness only requires a change in the chosen mate, the increase in reproductive investment needs physiological changes that enable this to happen, as seen in the late breeding season (Passos et al., 2021). As current reproduction trades-off with survival and future reproduction (Edward & Chapman, 2011; Stearns, 1989), the increase in reproductive investment could have a higher cost than the reduction of choosiness. In this scenario, the cues required to elicit changes in choosiness might be less exigent than the ones needed to do it in reproductive effort. It was proposed that the increase in reproductive investment depends on the whole context of individuals (including their internal state and the external environment), rather than in only one cue (Duffield et al., 2017). In that case, the solely decrease of water depth might have not been a signal complex enough to cause the increase in reproductive effort, but it could be an appropriate cue to reduce choosiness. Second, there could be differences between females and males in their behavioural plasticity. While mating depends on female acceptance, courtship and aggression do not. If behavioural plasticity is higher in females than males, desiccation risk may influence reproductive isolation but not reproductive effort, as most of the behaviours considered in this study are performed by males. However, even though there is some evidence of sex differences in behavioural plasticity, it is not clear how widespread they are (Fox et al., 2019; Hangartner et al., 2022).

When changes in environmental factors promote the reduction of choosiness but they do not reduce motivation to mate, hybridization could increase (Rosenthal, 2017). In response to reduced survival expectancy, *Garcialebias* decrease their choosiness and increase their sexual behaviour. Therefore, hybridization increases as a consequence of the reduction in survival expectancy. It is hard to predict how increased hybridization impacts natural populations since the outcome depends on several factors, including the hybrid fitness (Irwin & Schluter, 2022). Although hybridization could cause the collapse of parental populations (Irwin & Schluter, 2022) or hybrid speciation (Abbott et al., 2013; Abbott et al., 2010; Mallet, 2007), it could also reach a stable equilibrium (Irwin & Schluter, 2022; Servedio & Hermisson, 2020). In fact, when choosiness depends on the context, species might reach a partial reproductive isolation that stabilizes over time (Servedio & Hermisson, 2020). Hybridization can be a choice because it is better to hybridize than not to mate at all, but it could have another upsides (Pfennig, 2021). The temporary ponds that *Garcialebias* inhabit are usually isolated from each other, with little migration between populations (García et al., 2016). Thus, in this scenario, hybridization could be an advantage, helping *Garcialebias* to diminish the impacts of genetic drift and endogamy in their habitats (Garcia et al., 2020).

Our results show that reproductive isolation between *G. reicherti* and *G. charrua* varies depending on their survival expectancy, which causes reproductive isolation to change over a very short period of time. Many previous studies have examined how reproductive isolation varies in the geographic space (e.g. Sato et al., 2016; Yang et al., 2016; Yukilevich & Peterson, 2019), and over long periods of time (Kulmuni et al., 2020; Seehausen et al., 2014), however, less is known about how reproductive isolation varies over short periods of time. Our study highlights the temporal dynamics of reproductive isolation, an aspect that has been pointed out as necessary to understand hybridization and speciation (Schumer et al., 2017). Because of their particular lifecycle and their ability to cope with it by adjusting their physiology and behaviour, *Garcialebias* are great models to investigate the dynamics of hybridization and its evolutive consequences.

### Data Availability

Data set is available on figshare: https://figshare.com/s/831e280ad839abf5ec18

